# Distinguishing fine structure and summary representation of sound textures from neural activity

**DOI:** 10.1101/2022.03.17.484757

**Authors:** Martina Berto, Emiliano Ricciardi, Pietro Pietrini, Nathan Weisz, Davide Bottari

## Abstract

The auditory system relies on both local and summary representations; acoustic local features exceeding system constraints are compacted into a set of summary statistics. Such compression is pivotal for sound-object recognition. Here, we assessed whether computations subtending local and statistical representations of sounds could be distinguished at the neural level. A computational auditory model was employed to extract auditory statistics from natural sound textures (i.e., fire, rain) and to generate synthetic exemplars where local and statistical properties were controlled. Twenty-four human participants were passively exposed to auditory streams while the EEG was recorded. Each stream could consist of short, medium, or long sounds to vary the amount of acoustic information. Short and long sounds were expected to engage local or summary statistics representations, respectively. Data revealed a clear dissociation. Compared to summary-based ones, auditory-evoked responses based on local information were selectively greater in magnitude in short sounds. Opposite patterns emerged for longer sounds. Neural oscillations revealed that local features and summary statistics rely on neural activity occurring at different temporal scales, faster (beta) or slower (theta-alpha). These dissociations emerged automatically without explicit engagement in a discrimination task. Overall, this study demonstrates that the auditory system developed distinct coding mechanisms to discriminate changes in the acoustic environment based on fine structure and summary representations.

**SIGNIFICANCE STATEMENT:** Prior to this study, it was unknown whether we could measure auditory discrimination based on local temporal features or spectrotemporal statistics properties of sounds from brain responses. Results show that the two auditory modes of sound discrimination (local and summary statistics) are automatically attuned to the temporal resolution (high or low) at which a change has occurred. In line with the temporal resolutions of auditory statistics, faster or slower neural oscillations (temporal scales) code sound changes based on local or summary representations. These findings expand our knowledge of some fundamental mechanisms underlying the function of the auditory system.

## INTRODUCTION

The human auditory system can discriminate sounds at both high and low temporal resolutions (McAdams, 1993; Griffiths, 2001). The processing of fine temporal details relies on extracting and retaining local acoustic features (on the order of a few milliseconds) to detect transient changes over time (Plomp, 1964; McDermott, Schemitsch, and Simoncelli, 2013; Dau, Kollmeier, and Kohlrausch, 1997). These temporal variations characterize different sound objects and help the system discern among acoustic sources. However, environmental inputs typically comprise long-lasting sounds in which the number of local features to be retained exceeds the sensory storage capacity. For this reason, the system may need to condense information into more compact representations to discriminate sounds over more extended periods (McDermott, Schemitsch, and Simoncelli, 2013). As the duration of the entering sounds increases, summary representations are built upon fine-grained acoustic features to condense information into a more compact and retainable structure (Yabe et al., 1998). The processing of summary representations allows abstraction from local acoustic features and prompt sound categorization (McDermott and Simoncelli, 2011; McDermott, Schemitsch, and Simoncelli, 2013).

For sounds characterized by a constant repetition of similar events over time (such as sound textures, e.g., rain, fire, typewriting; Saint-Arnaud and Popat, 1995), this form of compression consists of a set of auditory statistics comprising averages over time of acoustic amplitude modulations at different frequencies (McDermott and Simoncelli, 2011; Figure 1A).

**Figure 1.**
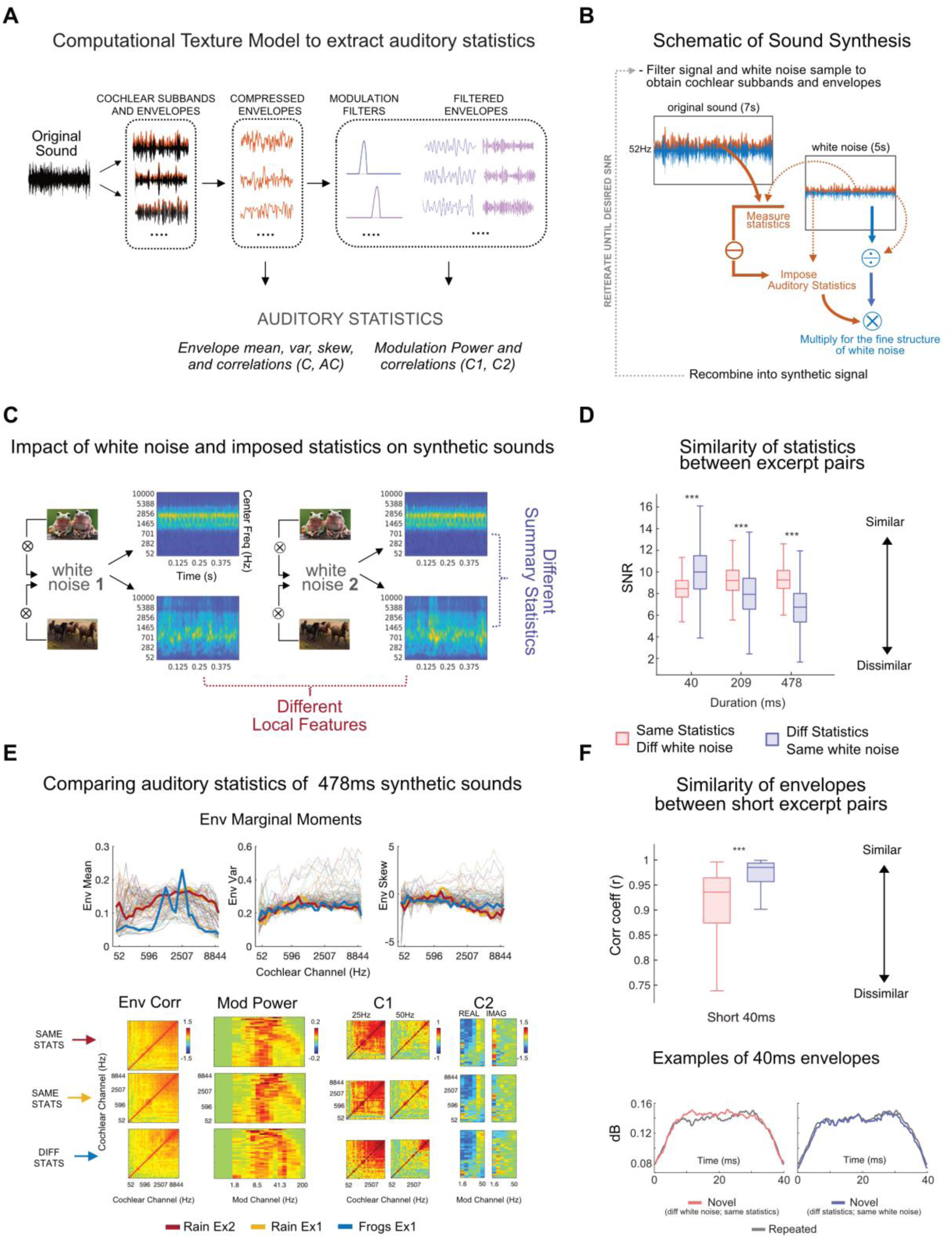
Experimental stimuli. (A) Computational Texture Model to extract auditory statistics. An original recording of a natural sound texture is passed through the auditory texture model (the list of presented sound textures is available as Extended Data, Figure 1-2). The model provides a mathematical formulation of the auditory system’s computations (auditory statistics) to represent the sound object. The signal is filtered with 32 audio filters to extract analytic and envelope modulations for each cochlear sub-band. Envelopes are downsampled and multiplied by a compression factor. From the compressed envelopes, a first set of statistics is computed: marginal moments (including envelope mean, variance, and skewness), autocorrelation between temporal intervals, and cross-band correlations. Compressed envelopes are then filtered with 20 modulation filters. The remaining statistics are extracted from the filtered envelopes: modulation power and cross-band correlations between envelopes filtered with the same modulation filter (C1) and between the same envelope filtered through different filters (C2). (B) Schematic of Sound Synthesis. The white-noise sample is filtered through the auditory model (McDermott and Simoncelli, 2011) to extract its cochlear envelopes, which are then subtracted from those obtained from the original sound texture. The average statistics from the original sound textures are then imposed on the subtracted white noise envelopes. The outcome is multiplied by the fine structure of the white noise sample to preserve its local acoustic distribution (e.g., temporal structure). The result is recombined in the synthetic signal, reiterating the procedure until a desired SNR of 20-dB is reached. (C) Impact of white noise sample and imposed statistics on synthetic sounds. Two different sets of statistics are extracted from two sound textures: “frogs” and “horse trotting”. Each set of values is imposed on two different random white noise samples. When the same statistics are imposed on different white noise samples, the outcomes are two synthetic exemplars of the same sound texture. These exemplars will have the same summary statistical representation but will diverge in their local features as the original input sound will influence them. When different statistics are imposed on the same white noise sample, the results are two synthetic exemplars that will diverge in their overall summary statistics and be perceptually associated with different sound objects. The cochleograms of the 0.5 s synthetic exemplars are displayed. (D) Similarity of statistics between excerpt pairs. Couples of sound excerpts presented in the study (repeated and novel; see Figure 2A for the experimental protocol) could be derived from different white noise samples to which we imposed the same statistics (in coral) or from the same white noise sample with different statistics (in blue). The summary statistics similarity between these couples of synthetic excerpts was computed by averaging the SNRs between statistics of repeated and novel sounds, measured separately for each statistical class. Boxplots show the averaged SNRs at three sound durations of interest (short, 40ms; medium, 209ms; long, 478ms). When sounds were short (40ms), statistical values were more similar for sounds derived from the same white noise samples (in blue) compared to different ones (in coral), even when including different original statistics. As duration increased (209, 478ms), statistics progressively converged to their original values and were more dissimilar for sounds with different generative statistics (blue) than for sounds including the same statistics (coral), irrespectively of original white noise sample. *** p < 0.001 (E) Comparing auditory statistics of 478ms synthetic sounds. Envelope marginal moments (mean, skewness, and variance) of all sound textures are displayed, while highlighted are those from three sound excerpts selected randomly; two have the same imposed auditory statistics (in red and yellow), and one has different statistics (in blue). In the bottom row, the remaining statistics are displayed (envelope correlation, modulation power, C1, and C2). The similarity between statistical values is higher when the sounds come from the same original texture. (F) Similarity between envelope pairs of short sounds. In the top panel, boxplots represent the correlation coefficients (r) measured between broadband envelopes for each pair of 40ms sound excerpts (repeated and novel; n= 6912) divided according to experiment (Local Features or Summary Statistics). Amplitude modulations of brief excerpts are significantly more similar when sound pairs originate from the same white noise sample (Summary Statistics experiment) than when they do not (as in the Local Features experiment), disregarding their imposed generative statistics. ***p< 0.001. In the bottom panel, show examples of the 40ms broadband envelopes used for computing the correlation coefficients (r) above.

Computational approaches in auditory neuroscience allow the mathematical formalization of this set of auditory statistics (Figure 1A). The basic assumption is derived from information theories (Barlow, 1961) and suggests that if the brain represents sensory input with a set of measurements (statistics), any signal containing values matching those measurements will be perceived as the same.

Psychophysical experiments revealed that stimuli including the same summary statistics-but different local features-are easy to discriminate when they are short, but that as duration increases and summary representation takes over, they are progressively more challenging to tell apart (Berto et al., 2021; McDermott, Schemitsch, and Simoncelli, 2013). On the other hand, when sounds comprise different statistics, their perceived dissimilarity will increase with duration as their summary representations diverge (Berto et al., 2021; McDermott, Schemitsch, and Simoncelli, 2013). While some evidence exists in the animal model (see Zhai et al., 2020, for results in rabbits), the neural activity underpinning local features and summary statistics is unknown in humans. Moreover, previous behavioral studies required participants to attend to stimuli to perform a task actively. From this evidence alone, it thus remains unanswered whether discrimination based on local features and their summary statistics can occur despite the lack of an active task and can therefore occur automatically.

To fill these gaps, we used a validated computational auditory model (McDermott and Simoncelli, 2011) to extract auditory summary statistics from natural sounds and generate synthetic sounds that feature this same set of measurements (see Material and Methods; Figure 1A,B). With this approach, it is possible to impose the same set of statistics on different white noise samples that initially had different local structures (Figure 1B,C). By employing this synthesis approach, we could create sounds that differ at high temporal resolutions (e.g., local features) but are perceptually indistinguishable at lower ones (summary statistics) and vice versa (Figure 1C). We acquired EEG measurements in participants passively exposed to streams composed of triplets of sounds presented at a fast stimulation rate (2Hz). To ensure generalizability, sounds were randomly drawn from a large set of synthetic excerpts (see Material and Methods). Within each triplet, the first sound was repeated twice, while the third one was novel. Two experiments were designed (Figure 2A). (1) In Local Features, the novel and repeated sounds differed only in their local structures, as they were generated by imposing the same auditory statistics on different white noise samples; (2) in Summary Statistics, the novel sound was generated from the same white noise sample but differed from the repeated ones as it comprised a different set of auditory statistics. As summary statistics are expected to be relevant at increasing sound duration (McDermott, Schemitsch, and Simoncelli, 2013), sounds including the same statistics but originating from different input white noises will be easily distinguishable at short duration but not at long ones (Figure 1D). By contrast, sounds derived from the same white noise sample but including different summary statistics will have different statistical values when measured at long durations but more similar values when measured at short durations (Figure 1D). In fact, at short durations, statistics will be influenced by their similar temporal structure (see Figure 1F).

**Figure 2.**
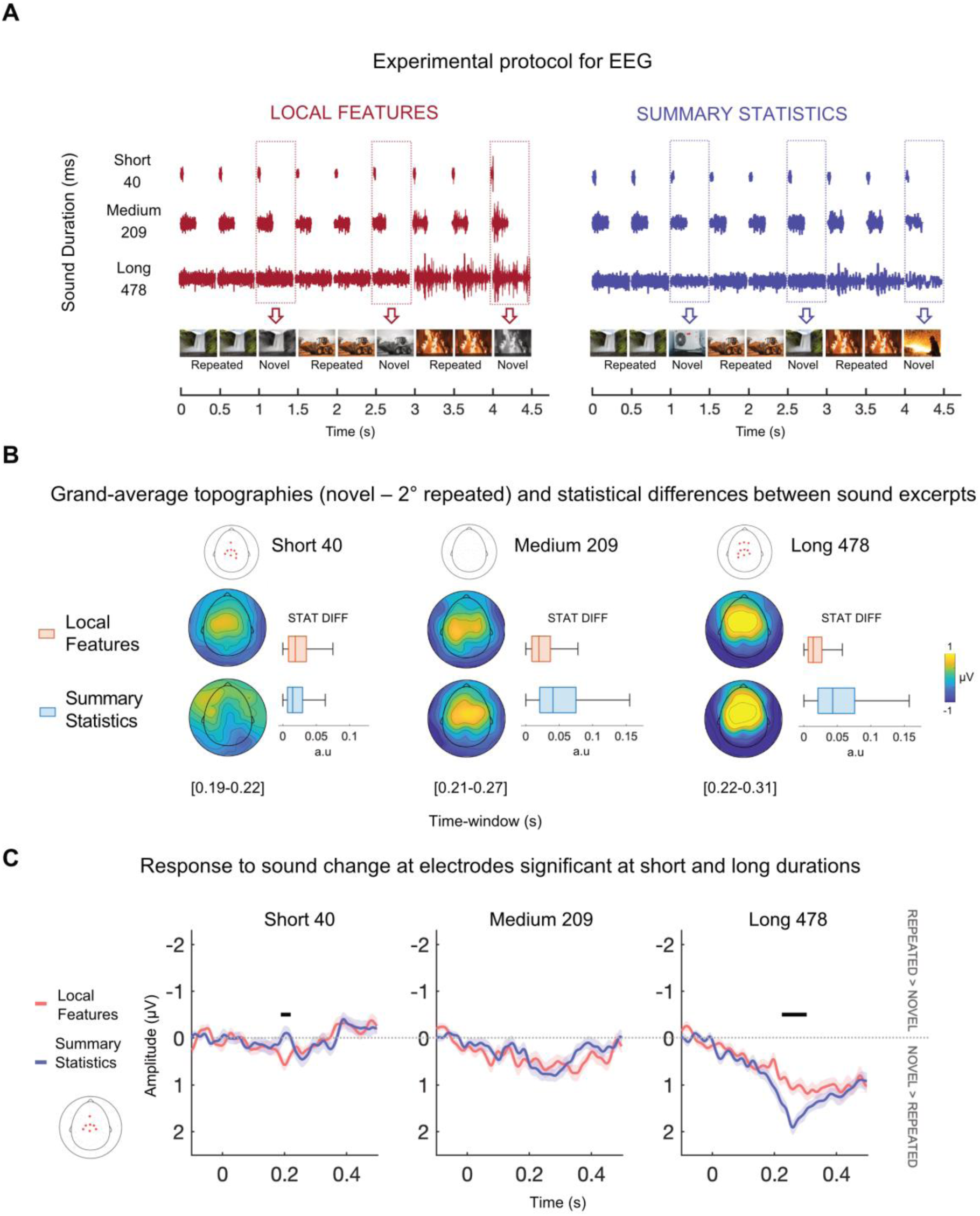
Experimental procedure and results of time domain analysis. (A) Experimental protocol for EEG. Triplets of sounds were presented at a fast rate (one sound every 500ms). Two sounds were identical (Repeated), while the third was different (Novel) and could vary in its local features (left) or summary statistics (right) depending on the experiment (Local Features or Summary Statistics). Three sound durations, equally spaced logarithmically (short, medium, and Long: 40, 209, and 478ms), were employed (in different sound streams) to tap into each auditory mode separately (local features vs. summary statistics processing). The list of presented sound textures is available as Extended Data, Figure 1-2. To ensure participants were attentive during the presentation, they performed an orthogonal task, consisting of pressing a button when an infrequent target (beep) appears. Performance accuracy was high in all experiments and durations and is displayed in Figure 1-1 in Extended Data. (B) Grand average topographies of the differential response associated with the sound change (novel sound minus repeated sound) at significant latencies for each experiment and duration. For each latency, electrodes associated with significant clusters are displayed above as red stars on the scalp. * p < 0.025. On the right side of the topographical maps, the boxplots represent objective differences between the novel and repeated sounds of all auditory statistics (averaged). The difference was computed between the statistics of sounds presented for each run, experiment, and duration and averaged across all participants. Within each duration, medians differed at the 5% significance level between experiments. Local Features > Summary Statistics at short (40) duration and Summary Statistics > Local Features for medium (209) and long (478) durations. The evoked response in the EEG agrees with the objective statistical difference measured from the sound excerpts. (C) Grand average electrical activity (negative values are plotted up) of the differential response (novel minus repeated) at significant electrodes (in red) for both short and long durations. Shaded regions show interpolated repeated error of the mean (SE) at each time point. Positive values indicate that the novel elicited a greater response than repeated. Results of cluster permutation are displayed as black bars extending through significant latencies. – p < 0.025. For visualizing the ERPs before subtraction (novel-repeated), see Extended Data, Figure 2-1.

Thus, to manipulate the extent of temporal and statistical similarity, we presented separate sound streams comprising stimuli of different lengths (either 40, 209, or 478ms; Figure 2A). First, we investigated auditory-evoked responses to uncover magnitude changes in neural activity associated with the two modes of representation. We predicted that short and long sounds would prompt larger auditory-discriminative responses for local features and summary statistics, respectively. Specifically, we hypothesized that since the amount of information (e.g., sound duration) impacts the statistical similarity of sound excerpts, distinct mechanisms are engaged in the processing of local features compared to summary statistics emerging over time. That is, in the case of short sounds, the brain may emphasize transient amplitude modulations (i.e., broadband envelope changes), while spectrotemporal statistics will become informative as sound size increases.

In line with this prediction, we expected brief local information to be encoded at a faster timescale (Panzeri et al., 2010) than summary statistics. That is, we expect the response pattern of the neuronal populations involved in processing local features to be encoded at higher frequency ranges and earlier latencies of neural oscillations compared with summary statistics. To this end, we investigated neural oscillations and assessed whether information measured at different temporal scales in the oscillatory pattern revealed specific fingerprints of discrimination based on local features and summary statistics.

## MATERIALS AND METHODS

### Participants

Twenty-four normal-hearing right-handed young adults (12 of either sex; mean age= 27.13 years, std= 2.83) participated in the experiment. All participants were healthy; they were fully informed of the scope of the experiment, signed written informed consent before testing, and received monetary compensation. The study was approved by the regional ethical committee, and the protocol adhered to the guidelines of the Declaration of Helsinki (2013).

### Sample size estimation

This sample size was estimated via simulations. We used the procedure described in Wang and Zhang (2021) and simulated a dataset with two conditions (Local Features and Summary Statistics) of Auditory Evoked Potentials data. First, we selected three electrodes of interest at central locations (E7, E65, E54). For the simulation, we chose a time window between 0.1 and 0.3s based on previous MMN studies (see Näätänen et al., 2007 for review). The amplitude values at the electrodes of interest for the two conditions were sampled from a bivariate normal distribution (within-subject design) where mean and standard deviation were chosen based on results of four pilot datasets (mean Local Features= 0.16; mean Summary Statistics= 0.56; std Local Features= 0.52; std Summary Statistics= 0.54).

We then ran a cluster-based permutation on simulated datasets to test whether any statistical cluster (t-values) exhibited a significant difference between the two conditions with an alpha level of 0.05. The procedure started with a sample size of 10 and increased in steps of one until it reached a power of 0.80. We ran 1000 simulations for each sample size and calculated the power as the proportion of the number of times significant clusters were found in these 1000 simulations. The simulation results showed that to obtain power above 0.8, a sample size of N= 24 was required.

The algorithm to perform such analyses can be downloaded from this link: https://osf.io/rmqhc/

### Stimuli

Synthetic sounds were generated using a previously validated computational auditory model of the periphery. The auditory model and synthesis toolbox are available at: http://mcdermottlab.mit.edu/downloads.html.

This auditory model emulates basic computations occurring in the cochlea and midbrain (McDermott and Simoncelli, 2011).

The signal (7s original recording of a sound texture, N=54; see Extended Data Table 1-2) was decomposed into 32 cochlear subbands using a set of gammatone filter banks with different central frequencies spaced on an ERB scale. Absolute values of the Hilbert transform for each subband were computed to extract the envelope modulation of each cochlear channel over time. Envelopes were then compressed to account for the nonlinear transformations performed by the cochlea. The first set of statistics was measured from the transformed envelopes: mean, skewness, variance, autocorrelation (within each cochlear channel), and cross-correlation (between channels). Additional filtering was applied to the envelopes to account for the modulatory response of the spectrotemporal receptive fields of neurons in the midbrain (Bacon and Wesley Grantham, 1989; Dau et al., 1997). Three additional statistics resulting from these operations could be derived: modulation power, C1, and C2 (respectively, the correlation between different envelopes filtered through the same modulation filter and the correlation between the same envelopes filtered by other modulation filters; Figure 1A). The resulting set of statistics extracted from the original recording of sound textures was imposed on four 5s white noise samples (Figure 1A, B, C). This allowed the generation of four different sound exemplars for each sound texture, which varied selectively in their local features but included similar long-term summary representations (Figure 1C). All synthetic exemplars featuring the same auditory statistics were perceptually very similar to the original sound texture from which they were derived, even when their input sounds (white noise) varied (Figure 1C-E). Synthetic sounds with the same imposed auditory statistics represent different exemplars of the same sound texture with the same summary statistics but a different fine-grained structure. This is because, in the synthesis procedure, the imposed statistics are combined with the fine structure of the original white noise sample (Figure 1B).

Importantly, to create experimental stimuli, all four 5s synthetic exemplars were cut from the beginning to the end into excerpts of different lengths, either short (40ms), medium (209ms) or long (478ms). These lengths were chosen based on results in previous behavioral investigations (Berto et al., 2021; McDermott, Schemitsch, and Simoncelli, 2013). Excerpts were equalized to the same root mean square amplitude (RMS= 0.1) and had a sampling rate of 20kHz. A 20ms ramp (half-hann window) was applied to each excerpt, 10ms at the beginning and 10ms at the end, to avoid edge artifacts (McDermott, Schemitsch, and Simoncelli, 2013). The stimuli used here were validated in a previous study (Berto et al., 2021) in which we replicated the original finding (McDermott, Schemitsch, and Simoncelli, 2013). The experimental stimuli presented for each run were randomly drawn from all available excerpts according to the experiment requests (see below).

### Procedure

Participants were tested in a sound-isolation booth. After reading instructions on a monitor, they listened to the sounds in the absence of retinal input (participants were blindfolded to prevent visual input).

For each run of the experimental session, a sound sequence lasting 108s was presented. The series contained triplets of sounds (n = 216) presented one after the other to form an almost continuous sound stream, in which sound onsets occurred every 500ms (Figure 2A). Within each sequence, all sounds had the same duration (either 40, 209, or 478ms).

Two experiments were implemented: (1) In Local Features, two different 5s synthetic exemplars of the same sound texture were selected (out of the four we had created); the combination of selected pair of exemplars vary randomly across triplets (e.g., first and second; second and fourth, and so on). These two exemplars were cut into brief excerpts of either 40, 209, or 478ms. According to the selected duration (which was different for each sequence), two excerpts (one for each exemplar) were chosen from among the available ones. The two excerpts had the same starting point (in seconds) from the onset of the 5s exemplar. The first sound excerpt was repeated twice, and afterward, the other was presented as the third element in the triplet.

Thus, two sounds within a triplet were identical (repeated), while the third one (novel) comprised different local features but converging summary statistics; in other words, repeated and novel sounds had the same generative statistics (both could be, e.g., waterfall) but different acoustic local features (Figure 2A, left panel; Extended Data Table 1-2, column 1). (2) In Summary Statistics, sound textures were coupled according to their perceived similarity (McDermott, Schemitsch, and Simoncelli, 2013; see Extended Data Table 1-2, columns 1 and 2). For the textures in column 1, one out of the four 5s synthetic exemplars was selected and cut into excerpts of the required duration (40, 209, or 478ms); one of such excerpts was picked randomly. The same was done for the coupled texture, ensuring that both were derived from the same white noise sample and that both drawn excerpts had the same starting point in seconds. Thus, we ensured that the sounds came from the same segment of the original input noise sample and varied only for their imposed statistics. Again, the first excerpt was repeated twice, while the other was used as the last sound in the triplet. The novel sound thus deviated from the other two (repeated) in its auditory statistics, as it was a segment of an exemplar of a different sound texture. This means the novel sound was a different sound object (e.g., the repeated sounds might be waterfall excerpts and the novel one air conditioner; see Figure 2A, right panel). However, since both originated from the same segment of the same input white noise sample, their temporal structure (i.e., broadband envelope) measured at high resolution (that is, in brief excerpts) was expected to be more similar in Summary Statistics than in the Local Features experiment. This was indeed the case (see Figure 1F) and would affect the similarity of statistics measured from short (but not long) sound excerpts (Figure 1D).

To ensure generalizability, the sound textures were different across triplets, so the statistical similarity between repeated and novel sounds was kept constant within an experiment while presenting different types of stationary sound objects.

Discriminative responses emerging from the contrast between the novel and repeated sounds did not depend on specific properties (e.g., a change in frequency between a particular type of sound category) but only on their local or statistical changes.

In both experiments, the order of the triplets was shuffled for each participant and run. Moreover, excerpts were selected randomly from among those that shared the required characteristics, so not only the presentation order but also stimuli per se were always different across participants.

A total of six conditions were employed: two experiments (Local Features and Summary Statistics) for three sound durations (40, 209, 478ms). Note that for each sound texture, we synthesized only four exemplars that we cut into excerpts of different sound durations (short, medium, or long). This means that within one experiment, the presented excerpts belonged to the same pool of synthetic sounds, and only their duration changed, not their properties. Thus, any dissociation between experiments (Local Features or Summary Statistics) according to sound duration would indicate that the processing of either local features or summary statistics strictly depends on the amount of information presented.

Two sequences/runs per condition (Experiment * Duration) were presented for a total of twelve runs. The order of runs was randomized across participants, and short breaks were taken between runs. In a sound stream, excerpts were presented in triplets, with the repeated one presented twice, followed by the novel one. Keeping the number of repeated sounds constant allowed to control for the effects that differences in their number could have on the brain response (e.g., standard formation, the effect by which the number of repeated stimuli influence the response to the deviant element; see Sussman and Gumeyuk, 2005); moreover, it allowed to keep the duration of the streams constant while manipulating the amount of information they encompass (e.g., the size of each sound excepts). On the other hand, by keeping the novel position fixed (as the third element of the triplet), we controlled for between-experiment differences in expectancy effects (e.g., some novel sounds could be more predictable than others at specific durations or based on their intrinsic properties) and, more importantly, we ensured that the novel sound varied from the repeated ones only for its generative statistics (same in Local Features and different in Summary Statistics) and original fine structure (different in Local Features and same in Summary Statistics).

Since the interstimulus gap always depended on sound duration (sound onset was kept constant at every 500ms), comparisons were assessed between experiments and within the duration.

Participants had to listen to the sound stream but were asked to perform an orthogonal task consisting of pressing a button when a beep sound was heard. The beep was a pure tone higher in pitch and intensity than the sound-texture stream. The pure tone was 50ms in length, had a frequency of 2200Hz, an amplitude of 50dB, a sampling rate of 20kHz, and an RMS of 5. The beeps randomly occurred during the stimulation period. The number of beeps varied randomly across runs from 0 to 3. Detection was considered valid when the participant pressed the key within an arbitrary window of 3s from beep occurrence.

### Similarity of summary statistics as a function of sound duration

In order to assess the impact of sound duration on the statistical similarity between pairs of excerpts, we extracted statistical values from all the pairs of excerpts (repeated and novel) presented in the experiment to all participants and in all runs (n= 20736; note that stimuli would appear more than once, as we adhered to the exact sound sequences presented to participants). That is, for each synthetic excerpt pair, we extracted the set of summary statistics (envelope mean, skewness, variance, and cross-band correlation; modulation power, C1, and C2) through the auditory texture model (Figure 1A; McDermott and Simoncelli, 2011). To assess similarity between summary statistic of repeated and novel sounds, we used a similar procedure to the one employed during sound synthesis to evaluates the quality of the output. This procedure consists of computing the signal-to-noise ratio (SNR) between statistic classes measured from the synthetic signal and the original sound texture (McDermott and Simoncelli, 2011). Firstly, we computed the total squared error *ε* of statistics measured from repeated sounds and the corresponding novel sound at each cochlear channel *k* (n=32) as follow:

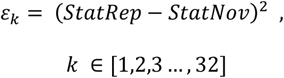

where *StatRep* is a statistic class (i.e., envelope mean, variance, or modulation power) measured from a repeated sound excerpt and *StatNov* is the same statistic class measured from the corresponding novel sound in the triplet. Note that for statistic classes that had more than the one dimension *k* (i.e., modulation power and correlations) the values across other dimensions (i.e., modulation bands) were summed prior to compute the error, as in McDermott and Simoncelli (2011).

Secondly, we calculated the SNR for each statistic class by dividing the sum of the squared statistic values measured from the repeated sound by the squared error between repeated and novel sounds as follow:

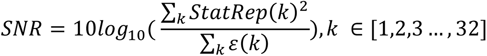

We computed one SNR for each statistic class (n=7) and then average their values to have one average SNR for each excerpt pair presented in each experiment and duration. Average SNRs are displayed in Figure 1D.

We then compared whether the average SNRs of sound excerpts were significantly different between experiment and within duration by performing non-parametric tests (Wilcoxon rank sum test). The results showed a clear dissociation according to sound duration. When sounds were short (40ms), the average SNR of statistics between repeated and novel sounds was higher in the Summary Statistics experiment (p < 0.001, mean=9.94; std=2.4) than in the Local Features one (mean= 8.34; std= 1.24). Namely, when sounds were short, statistical values were influenced by the white noise sample, thus sounds originated from the same seed had more similar values compared to when they originated from a different one, disregarding the generative statistics that were imposed. Thus, we expected larger neural discriminatory responses in Local Features experiment compared to the Summary Statistics one.

Conversely, at long duration (478ms), the average statistic SNR between repeated and novel sounds was more dissimilar in the Summary Statistics experiment (p < 0.001, mean= 6.73; std= 2.10) than in Local Features one (mean=9.31; std=1.23). At increasing sound duration, summary statistics were no longer influenced by the temporal structure of the original white noise sample as they converged to their original values. Based on this observation, we expected greater neural activation in response to Summary Statistics change compared to Local Features when sounds were long. The same pattern was observed for medium sound duration (209ms; p<0.001, mean Summary Statistics= 8.00; std= 2.21; mean Local Features=9.19; std=1.30), although there was a clear trend of decreasing average SNR with increasing sound duration in the Summary Statistics experiment (see Figure 1D).

Overall, this analysis showed that the statistical similarity measured from the presented sounds well predicted the brain response observed in the EEG.

### Similarity of temporal amplitude modulation in brief excerpts

The previous analysis showed higher statistical similarity measured at high (but not low) temporal resolutions from the excerpt pairs presented in the Summary Statistics experiment. To test the hypothesis that this effect depended on the original temporal structure of white noise samples (which will be more similar in the Summary Statistics experiment compared to the Local Features one), we conducted a similar correlation analysis for brief excerpts, but this time using as dependent variables the excerpts broadband amplitude modulations and disregarding their spectral density. Specifically, for every sound pair presented across participants, we used the auditory texture model (Figure 1A; McDermott and Simoncelli, 2011) to compute the cochleograms of all the 40ms excerpts presented in the study (n= 6912) and averaged them across frequency bands to extract their broadband envelopes. We then computed Pearson’s correlations between the envelopes of each excerpt pair (repeated and novel) to estimate their linear relationship (Figure 1F). The correlation coefficients (r) were transformed into Fisher-z scores for statistical comparison by t-tests. The results showed that the amplitude modulations over time between excerpt pairs were more correlated in the Summary Statistics experiment (mean = 2.35, std = 0.71) than in the Local Features experiment (mean = 1.63, std = 0.51, p-value <0.001). This result confirmed that regardless of their spectral density, the repeated and novel sounds in the Summary Statistics experiment shared more comparable temporal amplitude modulations than those in the Local Features experiment.

### EEG recording

Electroencephalography (EEG) was recorded from an EGI HydroCel Geodesic Sensor Net with 65 EEG channels and a Net Amps 400 amplifier (Electrical Geodesics, Inc., EGI, USA). The acquisition was obtained via EGI’s Net Station 5 software (Electrical Geodesics, Inc., EGI, USA). Central electrode E65 (Cz) was used as a reference. Four electrodes were located above the eyes and on the cheeks to capture eye movements. Electrode impedances were kept below 30 kΩ. The continuous EEG signal was recorded throughout the session with a sampling rate of 500Hz.

Experiment sounds were played from a stereo speaker (Bose Corporation, USA) positioned in front of the participant and at a 1m distance from the eyes; the sound level was kept constant across participants and runs (70dB). The experiment ran on MATLAB (R2018b; Natick, Massachusetts: The MathWorks Inc.); written instructions were displayed only at the beginning of the experimental session, via Psychtoolbox version 3 (Brainard and Vision, 1997; PTB-3; http://psychtoolbox.org/).

### EEG Data Analysis

#### Preprocessing

Data were preprocessed with a semi-automatic pipeline implemented in MATLAB (see Stropahl et al., 2018; Bottari et al., 2020). Preprocessing was performed using EEGLAB (Delorme and Makeig 2004; https://sccn.ucsd.edu/eeglab/index.php). Data were loaded, excluding electrode E65 (Cz), which was the reference channel of our EEG setup (thus consisting only of zero values).

A high-pass filter (windowed sinc FIR filter, cut-off frequency 0.1 Hz, and filter order 10000) was applied to the continuous signal to remove slow drifts and DC offset. A first segmentation in time was performed by epoching the signal according to the event onset. To avoid boundary artifacts, the signal was cut 2s before its onset event and until 2s after the end of the presentation (thus, from −2 to +114s) for each run. For each participant, epochs were merged in a single file containing only the parts of the signal referring to significant stimulation (thus excluding breaks between trials). Independent Component Analysis (ICA; Bell and Sejnowski, 1995; Jung et al., 2000a,b) was used to identify stereotypical artifacts. To improve ICA decomposition and reduce computational time, data were low-pass filtered (windowed sinc FIR filter, cut-off frequency 40Hz, filter order 50), downsampled to 250Hz, high-pass filtered (windowed sinc FIR filter, cut-off frequency 1Hz, filter order 500), and segmented into consecutive dummy epochs of 1s to spot non-stereotypical artifacts. Epochs with joint probability larger than three standard deviations were rejected (Bottari et al., 2020). PCA rank reduction was not applied before ICA to avoid compromising its quality and effectiveness (Artoni, Delorme, and Makeig, 2018).

For each subject, ICA weights were computed using the EEGLAB runica algorithm and then assigned to the corresponding original raw (unfiltered) dataset. Topographies for each component were plotted for visual inspection. Artifacts associated with eye movements and blinks were expected, and so a CORRMAP algorithm (Viola et al., 2009) was used to remove components associated with such artifacts semi-automatically. The automatic classification of components was performed using the EEGLAB plugin ICLabel (Pion-Tonachini, Kreutz-Delgrado, & Makeig, 2019).

Components representing eye movements and blinks were identified from their topographical map within the components ICLabel marked as ‘Eye’ with a percentage above 95%. Among these components, those with the highest rankings were selected from a single dataset and used as templates (one for eye movements and one for blinks). CORRMAP algorithm clusters ICA components with similar topography across all datasets to highlight the similarity between the IC template and all the other ICs. A correlation of the ICA inverse weights was computed, and similarity was allocated with a threshold criterion of correlation coefficient being equal to or greater than 0.8 (default value of CORRMAP; Viola et al., 2009). For all participants, on average, 1.92 components were removed (std= 0.88; range= 0-4).

Bad channels were interpolated after visually inspecting the scroll of the entire signal and power spectral density for each electrode. On average, 3.75 (range= 1-8; std= 2.21) channels were interpolated. The interpolation of noisy channels was performed via spherical interpolation implemented in EEGLAB.

Finally, the reference channel (Cz) was reintroduced in the EEG data of each participant, and the datasets were re-referenced to the average across all channels.

### Time domain analysis

This analysis was performed to extract auditory evoked potentials and uncover phase-locked magnitude changes associated with the two modes of sound representation (Local Features or Summary Statistics).

Pre-processed data were low-pass filtered (windowed sinc FIR filter, cut-off frequency= 40Hz, filter order= 50). Additionally, detrend was applied by filtering the data above 0.5Hz (windowed sinc FIR filter, cut-off frequency= 0.5Hz, filter order= 2000). Consecutive epochs (from −0.1 to 0.5s) were generated, including segments of either the novel sounds or the repeated one (the second) of the triplets for each participant and condition. Data were baseline corrected using the −0.1 to 0s pre-stimulus period. Specifically, we averaged all the time points from −100 to 0ms before the onset of each stimulus (either novel or repeated) and subtracted that value from post-stimulus activity (Luck, 2014). Joint probability was used to prune non-stereotypical artifacts (i.e., sudden increment of muscular activation); the rejection threshold was four standard deviations (Stropahl et al., 2018). For novel sounds, on average, 16.58 epochs per participant were removed (std=5.42; range 5-30) out of the 144 concatenated epochs that each Experiment * Duration comprised; for repeated sounds, on average, 16.15 epochs were removed (std= 5.11; range 5-29), again out of 144 trials per condition. Data was converted from EEGLAB to FieldTrip (Oostenveld, Fries, Maris, and Schoffelen, 2011; http://fieldtriptoolbox.org). Grand averages across participants were computed for each experiment, duration, and stimulus type (repeated or novel). Data across trials were averaged, generating Auditory Evoked Potentials (Figure 2-1 in Extended Data).

For each triplet, we subtracted from the evoked response to the novel sound the one to the preceding repeated one. Since all stimuli in the triplets (repeated and novel) were never the same across runs and participants, the subtraction was performed to ensure that neural responses were not driven by idiosyncratic differences in the stimuli that were presented in that specific run, but by the statistical difference between novel and repeated ones. Moreover, subtracting the response to the repeated sound from the one to the novel sound allowed us to isolate within-triplet differences from those between triplets. That is, since the first sound is repeated twice, the response to the second repetition is not independent of the brain activity elicited by the first one and likely incorporates a suppression mechanism to being exposed to the same stimulus twice. In the same vein, the subtraction metrics represented the relative distance between being exposed to the same sound as opposed to hearing a new one. Finally, the fact that in the two experiments, novel and repeated sounds varied for selective properties (either local features or summary statistics) allowed us to address how a deviation in fine temporal details or global statistics altered the response to sound change.

A nonparametric permutation test was performed between experiments (Local Features vs. Summary Statistics) for each duration (short, medium, and long), employing the subtracted auditory responses between the novel and repeated sounds. The permutation test was performed under the null hypothesis that probability distributions across condition-specific averages were identical across experiments.

The cluster-based permutation approach is a nonparametric test that has the advantage of solving the multiple comparison problem of multidimensional data in which you must control several variables, such as time, space, frequencies, and experimental conditions (Maris and Oostenveld, 2007).

Notably, statistical analyses between experiments were performed only within each duration to avoid possible confounds associated with refractoriness effects due to different interstimulus intervals (ISI) at long and short durations.

Thus, the contrasts of interest were: (1) Local Features short vs. Summary Statistics short; (2) Local Features medium vs. Summary Statistics medium; (3) Local Features long vs. Summary Statistics long.

A series of cluster-based permutation tests (Maris and Oostenveld, 2007; cluster alpha threshold of 0.05 (two-tailed, accounting for positive and negative clusters); 10000 permutations; minimum neighboring channels = 2) was performed. Cluster-based analyses were performed within a pool of central channels (according to EGI system, channels: E3, E4, E6, E7, E9, E16, E21, E41, E51, E54, E65); we selected the channels that better characterized the response to the second repeated sound, and which corresponded to the 11 central sensors we used in the analysis (e.g., see the topography in Extended Data, Figure 2-1). By pre-selecting this smaller number of central channels (whose response likely originates from auditory sources), we avoided including noisy channels in the model. Statistics were run for all samples from 0 to 0.5s. We expected novel sounds to elicit larger responses than repeated sounds.

### Time-Frequency analysis

Following the differences in magnitude changes observed between experiments for long and short durations, we performed data decomposition in the time-frequency domain to test whether sound changes at a high temporal resolution (local features in short sounds) were encoded at faster timescales compared to those occurring at a low temporal resolution (summary statistics in long sounds). We investigated frequencies below 40Hz, which have been associated with auditory processing in studies including both humans and animals (for review, see Gourevic et al., 2020). Specifically, several studies have marked the relevance of lower (theta, alpha) and higher (beta) frequency bands concerning auditory feature integration (e.g., VanRullen, 2016; Teng et al., 2018) and detection of deviant sounds (e.g., Fujioka et al., 2012; Snyder and Large, 2005). Preprocessed data were low-pass filtered to 100Hz (windowed sinc FIR filter, cut-off frequency= 100Hz, filter order= 20) to attenuate high frequencies and high-pass filtered at 0.5Hz (as with time-domain data). Data were epoched into segments from −0.5 to 1sec from stimulus onset: the second repeated or the novel. Joint probability was used to remove bad segments with a threshold of 4 standard deviations. On average, 11.96 epochs were removed for repeated sounds (range= 4-25; std= 4.28) and 11.58 for novel ones (range 4-26; std= 4.23). The resulting epoched datasets were converted to Fieldtrip for time-frequency analysis. We used complex Morlet wavelets to extract the power spectrum at each frequency of interest and time point. The frequencies spanned from 4 to 40Hz in steps of 2Hz; the time window for decomposition comprised latencies from −0.5 to 1s, around stimulus onset (either novel or repeated) in steps of 20ms.

Finally, the length of the wavelets (in cycles) increased linearly from 3 to 6.32 cycles with increasing frequency (depending on the number of frequencies to estimate; N=19). The signal was zero-padded at the beginning and end to ensure convolution with the central part of the window. The resulting power spectrum for each participant was averaged across trials.

Then, we performed a baseline correction to account for the power scaling (1/f). Unlike ERP analysis, baseline selection is a more sensible choice in time-frequency.

Therefore, it was crucial to choose a baseline whose position did not affect the results or over-boosted the effects. By using a stimulus-specific baseline as in the ERPs, for the novel sounds, we would be using as baseline the activity from a condition that, at least in some frequency ranges, is likely suppressed (the last 100ms of the response to the second repeated sound), while for the second repeated sound, we would be using as a baseline a segment in which activity is likely enhanced (as the first repeated sound includes between-triplet changes). Because of the nonlinearity of the baseline (to account for 1/f distribution), this will affect some frequencies more than others. When subtracting the power to the second repeated sound from the power measured for the novel sound, we would not be measuring the real dissimilarity between these responses, because the baseline correction would be unfair and so the relative power change. To account for this, we selected the same baseline for both the repeated and novel sounds, corresponding to the activity from −100 to 0ms before the second repeated sound. We decided to use a condition-averaged baseline (e.g., Cohen and Donner, 2013; Cohen and Cavanagh, 2011) to account for differences in the oscillatory tonic response as compared to the phasic one; since we are presenting a change always at the same rate, the activity could be phase-locked in time in a similar way across all the experiments, but the power at specific frequency bands could be higher in one experiment as compared to the other. If we used a condition-specific baseline, this effect would be masked because the activity would be corrected for the relative baseline measured during that stimulation stream. Therefore, we took the activity from 100ms prior to the onset of the second repeated sound for each experiment (Local Features or Summary Statistics) and averaged their power separately for each duration. As a baseline normalization method, we selected the relative change:

(pow(t)-bsl)/bsl

where pow is the total power at each sample (t) within the latencies of interest for repeated and novel grand-averaged trials, and bsl is the averaged baseline (across Experiment and time). The grand average of baseline-corrected power spectra of all participants was computed.

We investigated the neural activity underlying the discrimination of novel and repeated sounds across experiments for short and long durations. Thus, we first subtracted the power at repeated trials from that at novel trials and then used cluster-based permutation (Maris and Oostenveld, 2007) to investigate differences between neural responses to sound changes across experiments (Local Features vs. Summary Statistics) at each of the selected durations (short or long), at any latency (0 500ms) and across all (65) channels (minimum neighboring channels = 1). Following the inspection of power change between novel trials and repeated trials, oscillatory activity above 30Hz was not considered. We used the period of the oscillatory activity as an index of the temporal scale of the discriminative auditory processing, either slow, medium or fast. Since we did not have any apriori hypothesis concerning the contribution specific bands or ranges (e.g., from 9.5 to 16Hz) might have, we divided the power change into equally spaced frequency bands (each including 8 frequencies of interest, spaced in steps of 2Hz), creating a slow, medium, and fast oscillation range between 4 and 30Hz. These frequencies of interest included canonical theta, alpha, and beta oscillations (theta and alpha: 4-12Hz; low beta: 12-20Hz; high beta: 20-28Hz) but were unbiased by their canonical subdivision (for which theta would be 4-7Hz, alpha 8-12Hz, beta 13-25Hz and low gamma 25-40Hz). We instead hypothesized that the temporal scale of oscillation (from slower to higher) would encode the type of change that had occurred (local features vs. summary statistics). That is, depending on sound duration, we expected to detect different power modulations in response to changes in local features as compared to summary statistics at different timescales (frequency bands). Cluster permutation was performed separately for each frequency range (10000 permutations). The directionality of the test was based on results in the Auditory Evoked Responses (see Time-domain results) and on the specific frequency ranges: specifically, for a short duration, we expected power changes in higher frequencies in Local Features as compared to Summary Statistics. Conversely, at long duration, we expected greater power changes in the lower-frequency range in response to sound discrimination based on Summary Statistics compared with those based on Local Features. For the short duration, we thus expected: Local Features > Summary Statistics in the 4-12Hz range and Local Features < Summary Statistics in 12-20Hz and 20-28Hz. The opposite outcome was anticipated for the long duration: Summary Statistics > Local Features in the alpha-theta range; Summary Statistics < Local Features for beta bands (given the predefined directions of the effects, cluster alpha threshold was 0.05, one-tailed).

## RESULTS

### Behavioral Results

For each condition, the percentage of correct beep detections was above 90% (Local Features 40: mean= 0.99, std= 0.03; Local Features 209: mean= 0.99, std= 0.05; Local Features 478: mean= 1, std= 0; Summary Statistics 40: mean=0.99, std= 0.05; Summary Statistics 209: mean= 0.97, std=0.08; Summary Statistics 478: mean= 0.97, std= 0.11; Figure 1-1A, in Extended Data). We ran a two-way ANOVA for repeated measures with factors Experiment (2 levels, Local Features vs. Summary Statistics) and Duration (3 levels, 40, 209, and 478) to address whether experiment type and stimulus length had any impact on beep detection and participant attention to the task. No significant main effects were observed (Experiment, F(1,23)= 3.62, p =0.07, n2= 0.14; Duration, F(2,46)= 0.58, p= 0.56, n2= 0.3) or their interaction (Experiment*Duration, F(2,46)= 0.45, p= 0.64, n2= 0.2).

These behavioral results provide evidence that participants were attentive and responsive during sound presentation throughout the experiment and that attention to this orthogonal task was not influenced by the duration of the sound or experimental condition.

### Time domain results

By comparing Local Features vs. Summary Statistics separately for each sound duration, cluster permutation revealed a significant positive cluster, selectively for the short sound duration 40 (p < 0.02), lasting from 188 to 220ms after stimulus onset. Following the prediction, results revealed a greater auditory potential of Local Features compared to Summary Statistics for short duration. No significant positive cluster was found for the medium (209) and long (478) sound durations (all p > 0.39). Conversely, a significant negative cluster was found selectively for the long duration 478 (p < 0.001), lasting from 220 to 308ms after stimulus onset. These results indicate a greater response for Summary Statistics than Local features at long durations only. No differences emerged for short and medium sound durations (all ps >0.33).

Results clearly reveal double dissociations at the neural level based on stimulus length and mode of representation (Figure 2B,C). Findings support behavioral outcomes for which the processing of local features is favored for brief sound excerpts, while summary statistics are built at a slower temporal rate as information is accumulated (i.e., Berto et al., 2021; McDermott, Schemitsch, and Simoncelli, 2013). Going beyond past behavioral effects, our results clearly show that local and summary representations can emerge automatically from exposure to systematic sound changes. The neural response to an acoustic change depends on the similarity between local features and summary representations of sound excerpts. Summary statistics similarity can be manipulated as a function of sound duration, eliciting a dissociation in the magnitude of brain response that matches behavioral expectations.

### Time-Frequency Results

Since summary statistics emerge over time, we expected statistical variations to be encoded by slower oscillations than local feature changes. For such encoding, we expected power modulations at faster oscillations in response to local feature changes in short sounds and at slower oscillations in response to the emergence of a different set of summary statistics in long acoustic excerpts. To test this, we separated the power between 4 and 30Hz into three ranges, equally spaced: slow, 4-12Hz; medium, 16-20Hz; and fast, 20-28Hz. Then, we used a nonparametric permutation approach to address whether differences between Local Features and Summary Statistics emerged according to sound duration (short or long) within the three frequency ranges.

Results followed the predicted pattern. For the short sound duration, the analysis revealed a significant cluster between 100 and 220ms, in which sound change in Local Features elicited a greater decrease of power in the fastest oscillation range (20-28Hz; p< 0.05) compared to Summary Statistics (Figure 3A, left panel). This significant effect was located over left frontocentral and right posterior sensors (see Grand-average topography in Figure 3A, left). Conversely, for the long sound duration, we found a greater increase of power in the slow oscillation range for Summary Statistics compared to Local Features (4-12Hz; p < 0.03); the significant cluster consisted mainly of left frontocentral channels and bilateral posterior channels and spanned from 260 to 500ms (Figure 3A, right panel). No differences in power were found between Local Features and Summary Statistics for any sound duration in the medium frequency range (12-20Hz ranges, at any latency; all ps > 0.24). Overall, results revealed that when sound duration is short, neural oscillations at higher frequency bands (canonically corresponding to high-beta band) desynchronize more when the acoustic discrimination is driven solely by local features; vice-versa when sound duration is long, i.e., higher low-frequency oscillations (alpha and theta bands) are associated with stimulus changes based on different summary statistics (Figure 3B).

**Figure 3.**
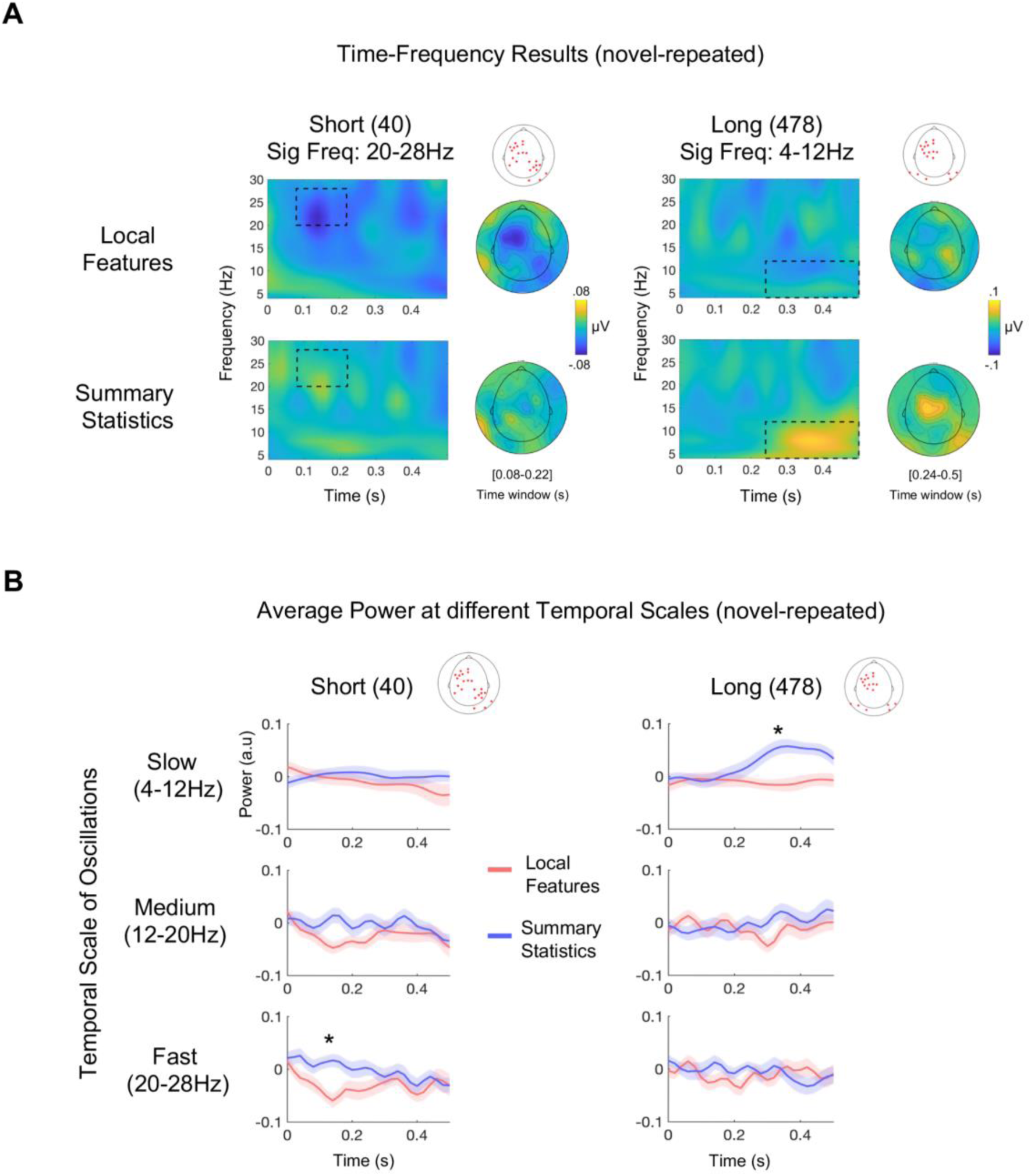
Results of time-frequency analysis. (A) Grand average difference (novel minus repeated) of total power for short and long sound durations in both experiments (Local Features and Summary Statistics) at significant channels. Rectangular regions comprise the latencies and frequency range in which power changes were significant between experiments after cluster-based permutation. Significant channels are marked as red stars over the sketch of a scalp (* p < 0.05). In the left panel, results for the short duration are displayed and show higher-power desynchronization in the 20-28Hz frequency range (high beta) for Local Features as compared to Summary Statistics. In the right panel, results for the long duration show higher 4-12Hz (alpha-theta) power synchronization for Summary Statistics as compared to Local Features. Grand-average topographical maps at significant latencies and frequency ranges are displayed next to the corresponding power-spectrum plots. (B) Average power difference between novel and repeated sounds for each range of frequency bands (Slow, Medium, and Fast), averaged across all significant channels, plotted at all latencies (from 0 to 0.5s). Significant channels are marked as red stars over the sketch of a scalp. Shaded regions show interpolated standard error of the mean (SE) at each time point. * p < 0.05.

Overall, these findings show that different temporal scales at the neural level underpin the discrimination of variant elements in the auditory environment based on the amount of information available and the type of sound change that has occurred.

Notably, beta desynchronization for Local Features (short duration) peaks 100-150ms after stimulus onset, while the same effect in the time domain has a peak that builds up around 200ms. The opposite was found for Summary Statistics (long duration), in which theta-alpha synchronization starts about 40ms later than the effect observed in the time domain and is more sustained over time (i.e., it lasts the entire time window). These differences are indicative that the two measures capture at least partly different aspects of sound discrimination.

## DISCUSSION

The auditory system extracts information at high (local) and low (summary) temporal resolutions. Here, we assessed whether discriminative responses to local or summary representations could be measured at the neural level and whether they are encoded at different temporal scales (Panzeri et al., 2010). We employed a computational model (McDermott and Simoncelli, 2011) to synthetically create stimuli with the same summary statistics but different local features. We used these synthetic stimuli to present streams of triplets containing repeated and novel sounds that could vary in their local features or summary statistics.

Results in the time domain showed that when the sound duration was short, the magnitude of auditory potentials increased selectively for changes in local features. In contrast, when the sound duration was long, changes in auditory statistics elicited a higher response compared with changes in local features (Figure 2B, C). Thus, according to sound duration, we observed an opposite trend in the magnitude change of the evoked response. Note that for each sound texture, we manipulated the duration of the excerpts, and not their properties (we synthesized only 4 synthetic exemplars per sound texture, that we cut into smaller excerpts either 40, 209, or 478ms which were then randomly drawn in the experiments; see Material and Methods above). The dissociation observed between experiments according to sound duration is indicative that the processing of local features or summary statistics is strictly dependent on the amount of information presented. This trend perfectly matched expectations based on previous psychophysics evaluations (i.e., Berto et al., 2021; McDermott et al., 2013) despite the protocol was slightly different from the behavioral implementation. In the psychophysical version, the two experiments (Local and Summary) were substantially different from each other. One experiment, called Exemplar Discrimination, was the equivalent of the Local Features experiment in our protocol and contained two different sounds (since one was repeated twice). However, the other experiment, named Texture Discrimination, contained three different sound excerpts (two derived from the same white noise but with different imposed summary statistics; one derived from a different white noise with the same statistics). Different task demands justified this disparity.

Specifically, in the behavioral version, participants were given very clear instructions on which sound properties to pay attention to during each experiment (sound details or sound source, respectively) and even which sound to use for comparison (the middle one; McDermott et al., 2013). In this protocol, the sequences had the same structure in both experiments (two repeated sounds followed by a novel one), while the only difference was the generative statistics imposed on the novel sound compared to the repeated one (same in Local and different in Summary) or the white noise sample used to initialize the synthesis (different in Local and same in Summary). This allowed us to test for the automaticity of the processes and to measure distinct neural responses when the system is exposed to a similar or different set of statistics combined with the same or different local structure. Moreover, it permitted a fair comparison between experiments. Nonetheless, results went in the same direction in both the EEG and the behavioral evaluations, suggesting similar mechanisms are in place despite the lack of an explicit request to pay attention to specific sound properties.

Finally, analysis in the time-frequency domain revealed that neural activity at different temporal scales characterized discriminative responses to local features or summary statistics. Faster oscillations (in the beta range) were associated with discrimination based on local features, and slower oscillations (in the theta-alpha range) with changes based on summary statistics.

### Automaticity of Local Features and Summary Statistics Processing

Auditory responses to novel local features or summary statistics were associated with differences in magnitude that could be automatically detected. This finding confirms that the auditory system can attune its response to specific sound changes and expands seminal studies measuring the mismatch negativity (MMN) response (Näätänen et al., 1978; Tiitinen et al., 1994). MMN is the neural marker of a process by which the system “scans” for regularities in entering sounds and uses them as references to detect variations in the auditory scene (for reviews, see Näätänen et al., 2001, 2010). In our study, expectations that a change would occur in the third element of the triplet had a probability of 1 in each experiment (Local Features and Summary Statistics; Figure 2A). Thus, spurious expectancy or attentional effects cannot explain results. Coherently, the MMN response to a deviant sound is not affected by prior expectations that the novel element will occur (Rinne et al., 2001); rather, the auditory system automatically orients attention toward it. Here we highlighted another ability of the system. Beyond automatic orientation toward a relevant deviant sound, our results show that it is possible to categorize the acoustic change according to the representation (local or summary) and temporal resolution (high or low) at which it has occurred. Importantly, discriminative neural responses could be detected even if the task per se did not involve any discrimination or in-depth processing of either local features or summary statistics. In other words, the sound changes were processed even when irrelevant to the behavioral task participants were attending (rare beep detection), strongly suggesting that the entrainment to local or global acoustic change emerges automatically from exposure to regular changes in the environment and is strictly dependent on the amount of information presented.

Furthermore, the double dissociation we observed based on sound duration (with Local Features eliciting greater magnitude change than Summary Statistics for short sounds and vice-versa for long sounds) rules out the possibility of results being explained by a mere saliency effect (i.e., the fact that, in Summary Statistics, a different sound object was presented). Importantly, the main advantage of using synthetic sounds instead of natural recordings was to be able to control the summary statistics embedded in the sounds. That is, all sounds were random white noise samples to which we imposed the same (or a different set) of summary statistics. If the brain were not automatically encoding the summary statistics, we would not have been able to distinguish between Local Features and Summary Statistics experiments, especially at long duration, since all repeated and novel sounds differed for their local structure. Nor would it have been possible to detect a dissociation in the neural response according to sound duration.

This observation is further supported by the fact that results emerged despite sound objects between the triplets being continuously changing (the only fixed parameter was the expected similarity in local features or summary statistics between the novel and repeated sounds).

These findings can be generalized to a variety of sound textures (Figure 2A; see also Extended Data, Table 1-2) and the exact moment in which the summary percepts emerge likely depends on specific comparisons across sound objects (repeated and novel). In line with this, using many different sounds to create sound streams led to grand averaged signals associated with discrimination based on summary statistics with a rather spread-out shape (see Figure 2C, right).

Finally, it is important to notice that imposing different statistics on the same white noise leads to sounds with different long-term average spectra. Therefore, it is possible that magnitude differences in response to the Summary Statistics experiment, compared to Local Features, were driven by low-level spectrotemporal modulations rather than changes in higher-order statistics. However, if that was the case, we might have expected an effect already at medium duration (209ms), which was instead not present. Further experiments may be required to fully rule out this possible confound.

### Local features changes are encoded by fast oscillations

By comparing the difference in total power between novel and repeated sounds in the two experiments, we found that, for short sounds, the power between 20 and 28Hz decreased when a change in local features was detected, as compared to when summary statistics were changed. This desynchronization occurred between 80 and 200ms after stimulus onset (Figure 3A, B, left). Desynchronization of oscillatory activity is the decrease in power measured at specific frequency bands (generally alpha and beta ranges), which generally emerges following the onset of an event (Pfurtscheller and Lopes da Silva, 1999). It results from increased cellular excitability in thalamocortical circuits and generally reflects cortical withdrawal from the resting state to engage in a cognitive process (Pfurtscheller and Lopes da Silva, 1999).

The 20-28Hz band includes frequencies that are canonically attributed to high-beta oscillations. Changes in power synchronization in the beta range have been correlated with performance in tasks involving the detection of temporal or intensity deviations (Arnal et al., 2015; Herrmann et al., 2016). Overall, these findings suggest that, among other operations, brain activity in the high beta range could be engaged in the processing of low-level properties of a stimulus. Beta-band activity has also been investigated in the context of rhythmic perception. A disruption in beta power can be observed in non-rhythmic sequences or when an attended tone is omitted from a regular series (e.g., Fujioka et al., 2012). Interestingly, beta synchronization not only captures irregularities in a pattern but also reflects the type of change that has occurred. For instance, it has been shown that beta desynchronization was higher prior to the occurrence of a deviant sound whose pitch varied in a predictable way, as compared to an unpredictable variation. Accordingly, beta desynchronization has been proposed as a marker of predictive coding (Engel and Fries, 2010; Chang, Bosnyak, and Trainor, 2018).

In our model, stimuli could be derived from the same white noise sample or a different one (Figure 1C). In Local Features, the novel sound is derived from another white noise sample, as compared to the repeated sound on which we imposed the same summary statistics. Thus, with this synthesis approach, in terms of fine acoustic features, when sounds were short, novel sounds had a more different temporal structure (Figure 1F) and were statistically more dissimilar (Figure 1D, 2B) than their paired repeated one in the Local Features experiment as compared to Summary Statistics. Overall, these results suggest that, in the absence of enough information to build summary representations, faster oscillations are in charge of small, acoustic change detection to be used to discriminate sound excerpts.

### Slower oscillations are engaged in Summary Statistics processing

By comparing Local Features with Summary Statistics at long durations, we observed that the emergence of different auditory statistics in the novel sound, as compared to the previous, repeated one, elicited higher power at slower frequencies, compatible with canonical alpha-theta oscillations. This power synchronization emerged at relatively late latencies from stimulus onset (between 240 and 500ms; Figure 3A, B, right). and was not present when solely local features were driving sound change (as in the Local Features experiment). Provided that summary statistics can primarily be measured at increasing sound duration, we expected differences between long-duration stimuli being carried by relatively slower brain activity. However, statistical comparisons were performed within the sound duration; thus, if this effect was simply driven by the sounds being longer (478ms) rather than the processing of auditory statistics, we should not have observed a difference in alpha-theta synchronization between experiments.

Similarly, if the effect were driven by simply presenting a “more different” sound in the Summary Statistics experiment, as compared to Local Features one, then we would have seen an effect also for 209ms, which was not the case; similarly, we would not have been able to dissociate the effects based on sound duration. Finally, it is worth noting that the stimulation rate was kept constant across all tested durations (40, 209, and 478), meaning that we always presented one sound every half a second. This means that, disregarding the amount of information we presented, the change always occurred in a window of 1.5 seconds (with novel sound always occurring at a frequency of 0.667Hz). Therefore, the effect strictly depends on the amount of information we presented within this temporal window, rather than the time interval between sound excerpts.

A previous study investigated the temporal window of integration of sound textures, showing that it can extend for several seconds (McWalter and McDermott, 2018, 2019). In this study, we could not use stimuli longer than 500ms to maintain the 2Hz rhythmic stimulation pattern in all experiments. Thus, we could not address the integration effects of single sounds at longer durations. Interestingly, the integration window measured for sound textures is relatively long compared to the receptive fields of auditory neurons, whose response has been shown to be sustained for about a few hundred milliseconds (e.g., Miller et al., 2002). Previous evidence suggested the existence of an active chunking mechanism condensing entering acoustic information within a much longer temporal window, approximately 150-300ms (VanRullen, 2016; Riecke, Sacks, & Schroeder., 2015; Teng et al., 2018). Such integration length would be related to ongoing oscillatory cycles, specifically corresponding to the theta range (4-7Hz; Ghitza & Greenberg, 2009; Ghitza, 2012). Compatibly, a recent study showed that acoustic changes occurring around 200ms could explain the modulations of phase synchronization in theta (Teng et al., 2018).

Although there is no evidence that 200ms windows are relevant for texture perception (see McWalter and McDermott, 2018, 2019), our data show that brain activity already synchronizes 200ms after stimulus onset to the emergence of a novel set of auditory statistics. The integration window of sound texture defined by previous studies refers to the maximum duration within which the averaging of local information into summary statistics can occur (McWalter and McDermott, 2018). It is still unclear how this relates to the emergence of relevant percepts in the brain (i.e., sound object identity) in response to average statistics. The higher power synchronization in the theta-alpha range observed in response to sensory statistics might be interpreted as one of the possible neural mechanisms underlying the development of such abstract representations, which may lead to the perceptual understanding that a new sound object has occurred. This would explain why it happens when a different set of statistics is detected and not when only local features change while sound identity remains unchanged.

## CONCLUSION

Combining a computational synthesis approach with electrophysiology revealed distinct cortical representations associated with local and summary representations. We showed that different neural codes at faster and slower temporal scales are entrained to automatically detect changes in entering sounds based on summary statistics similarity emerging as a function of sound duration. These results promote using computational methods to appoint neural markers for basic auditory computation in fundamental and applied research. Furthermore, the automaticity of the protocol and the fast implementation allow the testing of different populations (including newborns, infants, children, and clinical patients) that do not have the resources to attend to complex tasks.

## DATA AVAILABILITY

Raw EEG data, analysis scripts, participants’ information, and sound excerpts employed in the experiment are available in an online repository at this link: https://data.mendeley.com/datasets/gx7cb7fnv4/1

## ACKNOWLEDGMENTS

The authors thank all the students who helped with recruiting participants and/or data collection: Nicolò Castellani, Irene Sanchez, Chiara Battaglini, and Dila Suay.

