## Extended Data for "Distinguishing fine structure and summary representation of sound textures from neural activity"

This document contains:

Extended Data Figure 1-1
Extended Data Figure 1-2 (Table)
Extended Data Figure 2-1

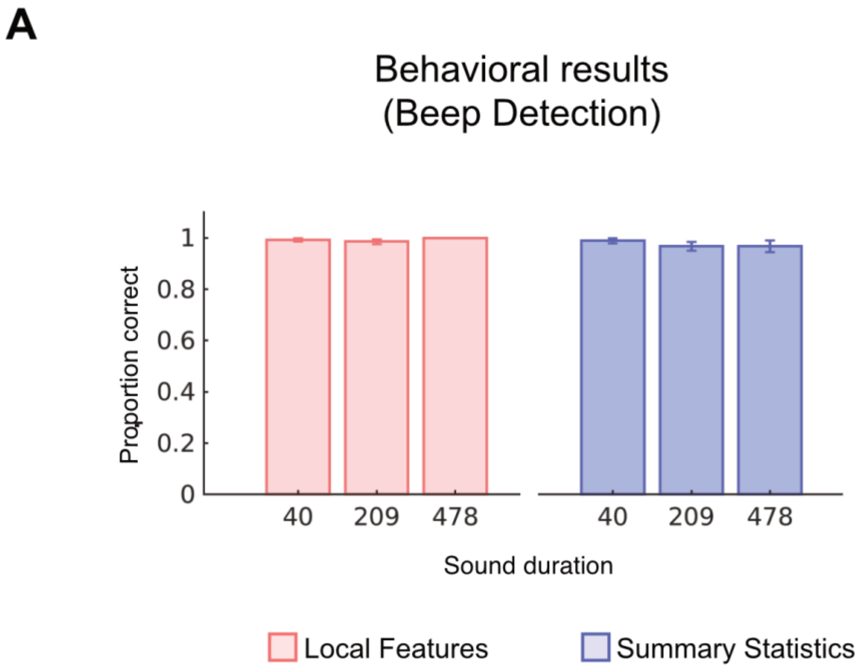


**Figure 1-1. Behavioral results. Related to Figure 2.** (A) The group-level average proportion of correct detections of beeps when presented. Bar plots represent average values of hits across all participants. Error bars represent the standard error of the mean (SE). No significant difference existed across conditions (all $p > 0.05$).

List of the sound textures employed in the experiments

| 1. | 2. |
|---|---|
| Applause large crowd | Applause big room |
| Applause 2 | Applause 1 |
| Bathroom sink | Bath being drawn |
| Bulldozer | Waterfall |
| Castanets 1 | Castanets 2 |
| Electric adding machine | Teletype city room |
| Fast running river | River running over shallows |
| Fire burning room 2 | Fire 3 |
| Fire forest inferno | Fire 3 |
| Frogs 4 | Frogs 3 |
| Frying bacon | Crunching cellophane |
| Heavy rain falling and dripping | Heavy rain on hard surface |
| Heavy rain on hard surface | Rain in woods2 |
| Horse trotting on cobblestones | Horse and buggy |
| Industrial machinery | Construction site ambience |
| Jungle rain | Rain in woods1 |
| Linotype | Teletype city room |
| Motorcycle idling | Idling boat |
| Pneumatic drills | Construction site ambience |
| Printing press | Construction site ambience |
| Radio static 2 | Radio static1 |
| Rain in woods 1 | Frogs 1 |
| Rain in woods 2 | Jungle rain |
| Rain | Rain in woods 1 |
| Rhythmic applause | Applause big room |
| River running over shallows | Applause big room |
| Shaking coins | Pouring coins1 |
| Ship anchor being raised | Pneumatic drills |
| Sparrows large, excited group | Birds in tropical forest |
| Stream near small waterfall | River running over shallows |
| Teletype city room | Teletype |
| Typewriter IBM electric | Typewriter manual |
| Water running into sink | Bathroom sink |
| Waterfall | Air conditioner |
| Frogs 3 | Frogs 1 |
| Metal lathe | Blender |
| Applause large crowd | Applause big room |

**Figure 1-2. List of Sound Textures. Related to Figure 1 and 2.** In Local Features Discrimination, for each sound texture in column 1, two synthetic exemplars of the sound texture were selected. One was presented twice (repeated) and the other was presented as the third element of the triplet (novel). In Summary Statistics Discrimination, sound textures were paired according to perceived similarity (McDermott, Schemitsch, and Simoncelli, 2013). For each sound texture in column 1, one synthetic exemplar was selected and presented twice. Then, an exemplar of the texture from the corresponding row in column 2 was selected and used as the third element of the triplet (novel).

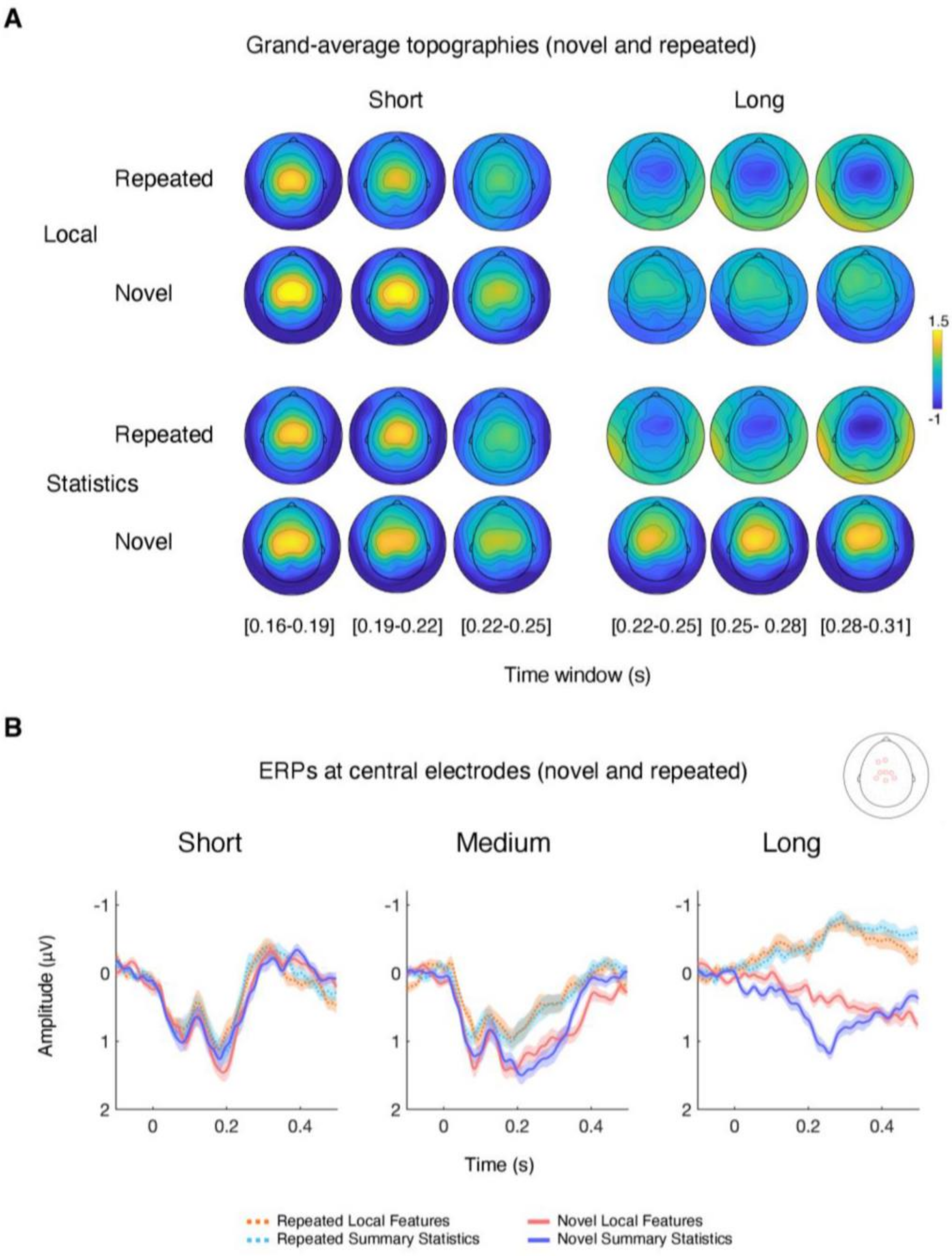


**Figure 2-1. Auditory Evoked response for repeated and novel sounds. Related to Figure 2.** (A) Grand-average topographies across participants of the responses to standard and oddball sounds for each experiment (Local and Global Discrimination), displayed for short and long durations (478) at latencies of interest. (B) Grand-average ERPs across participants of the average amplitude of the central channels displayed in the legend (red circles on the sketch of a scalp). ERPs are shown for both standard and oddball sounds for each experiment and duration. Shaded regions show interpolated standard error of the mean (SE) at each point.
